# Neural patterns associated with mixed valence feelings differ in consistency and predictability throughout the brain

**DOI:** 10.1101/2023.11.22.568316

**Authors:** Anthony G. Vaccaro, Helen Wu, Rishab Iyer, Shruti Shakthivel, Nina C. Christie, Antonio Damasio, Jonas Kaplan

## Abstract

Mixed valence feelings, defined by the simultaneous presence of positive and negative affect, remain understudied in neuroscience. We used functional magnetic resonance imaging to investigate subjects watching an animated short film known to induce mixed feelings. These same subjects were asked to identify the time periods during which they had experienced positive, negative, and mixed feelings. Using Hidden-Markov models, we found that activity in the insula, amygdala, nucleus accumbens, and anterior cingulate allowed us to predict the onset of new feeling states as determined by individual self-report. Further analyses found spatiotemporally unique and consistent neural patterns in the insular cortex for univalent states, but not for mixed valence states. However, ventromedial prefrontal cortex and anterior cingulate exhibited unique neural consistency for both univalent and mixed valence states. This study is the first to reveal direct evidence for a neurally consistent representation of mixed feelings in the central nervous system.

## Introduction

Mixed valence feelings (commonly referred to as “mixed feelings”), consist of simultaneously experiencing positive, negative, and generally conflicted feelings. Mixed feelings are usually thought of as a rarity but are actually common, and occur across cultures ^1, 2, 3, 4^. Events that trigger mixed feelings are often ascribed a significant personal meaning. ^5, 6, 7^. Despite their ubiquity, however, the topic of mixed feelings is largely absent from affective neuroscience research. One reason why mixed feelings are rarely studied may be their lack of fit with the dominant ways of thinking about emotion in neuroscience. In the constructionist approach, bipolar valence is an irreducible dimension of affect and consequently, positivity and negativity cannot simultaneously occur ^8, 9, 10^. In frameworks that view emotions as stemming from discrete functional biological states ^11^ the core affective systems and the traditionally defined emotion categories are generally conceived as solely positive or negative ^12, 13, 14^. When affective neuroscience research conducts experiments in service of finding evidence for or against these two theories, important topics such as mixed feelings tend to fall between the gaps ^15, 16^. Critically, the lack of research on mixed feelings limits our knowledge of how the neurobiology of valence actually works – an essential topic for understanding the biology of affective behaviors and subjective experiences. Of note, circuit-level studies in animals have suggested that there is flexible modularity in the limbic system for the representation of valence, where some groups of neurons interact in ways which allow bivalent representations, while others are more univalent or bipolar. Additionally, there is flexibility of these types of representations across contexts and types of modulation ^36, 37^. These findings suggest that concurrent positive and negative valence representation is be plausible, although subjective reports are necessary to provide direct evidence for a consistent neural counterpart to a ‘mixed’ feeling state.

In a previous theoretical paper, we advanced several hypotheses concerning the neurobiological foundations of mixed feelings ^16^. Our model posits that even when the neural correlates of emotions vacillate rapidly in the brainstem and subcortical limbic structures, subsequent integrative processes in the insula, result in a unified feeling state. With this model in mind, we may expect differences in the regions where state changes can accurately predict state changes during a film known to induce complex and mixed feelings. Specifically, insular cortex, ventromedial prefrontal cortex, and cingulate cortex may be predictive due to their roles in integrating various sources of affective information ^38, 39^. Furthermore, our framework predicts regional differences in whether mixed feelings are associated with a consistent neural state. If some functional circuitries encode positivity and negativity as mutually exclusive states ^40^, then the neural patterns when subjects are experiencing mixed feelings should be in flux. This would result in the regional patterns at different timepoints of experiencing mixed feelings, to be no more correlated with each other than they are with the regional patterns during the positive and negative states that they are switching between. On the other hand, when regions represent the feeling of mixed valence as an integrated and unique state ^41^, the regional pattern during mixed feelings should be correlated with other timepoints of itself than with these positive and negative states ^16^.

An additional problem with researching mixed feelings is the challenges they pose to our usual methods of research. Traditional emotion research in fMRI relies on the ability to have subjects feel similarly when experiencing the same stimuli. This increases our analytic power by allowing us to control for differences between stimuli in order to isolate the quality we are interested in, and by combining our data across subjects. But mixed feelings are not that easy to induce consistently across subjects, especially in a laboratory setting ^17, 18^, as they generally require contexts which allow for the contrast between the present view and another point in time^19^. Mixed feelings research can benefit from, and likely requires, using approaches and stimuli that allow the consideration of individual experiences rather than consistent responses across participants ^20, 21, 22^. This is why the recent trend of using more naturalistic types of stimuli, such as film, and finding creative ways to analyze them, is the perfect environment for research on mixed feelings ^23, 24^. Film stimuli are excellent for inducing mixed feelings because they are dynamic enough to generate the proper context for emotional conflict between the current moment and either the past or the future^25^. Viewers may even value the controlled ability to contemplate life that comes from bittersweet feelings in entertainment, more so than the purely pleasurable variety of feelings^6, 28, 29^. Mixed feelings induced by media may also facilitate coping with ambivalence. Engaging with narratives that induce such feelings can reduce delay-discounting ^30^ and induce greater reflection and acceptance of complex emotional issues ^31, 32^. In order to analyze the more dynamic, naturalistic stimuli that capture these complex emotions, we cannot rely on traditional univariate approaches.

Methods such as Hidden-Markov models are particularly suited to analyzing brain activity during film stimuli. With HMM, we assume that regional activity in the brain shifts over a number of distinct states, each with its own consistent and unique neural signature ^33^. With this type of analysis, we can test hypotheses about when state shifts occur in relation to the continuous stimuli, as well as regional-based hypotheses as to the features of the stimuli relevant to shifts in states ^12, 23^. HMM approaches have revealed that state changes in regions such as posterior medial cortex and prefrontal cortex align well with meaningful narrative events in film, whereas auditory and visual regions align instead with lower-level sensory features ^33, 34, 35^.

Recent studies have begun to apply HMM analyses to fMRI studies of affective experience. In one such study, states of the ventromedial prefrontal cortex were found to associate with affective states while watching a TV drama, though the timing of state onsets varied across subjects ^11^. The heterogeneity of these state change times highlights the importance of analyzing the neural dynamics of affect on an individual basis. However, this study assumed the same number of state changes for each participant, and did not link subject’s individual affective experience to their specific data. These two limitations, which were raised by the authors, are important and need to be tested for feasibility so as to allow affective neuroscience to study experiences as individual as mixed feelings. There is one recent fMRI study focused specifically on mixed feelings, investigating amusing but disgusting clips. The study found significant activity in the posterior cingulate associated with amusement mixed with disgust, but the specific patterns were not significantly different than those that occurred during disgust ^42^. This study focused on ambiguous social scenarios which may represent a different phenomenon from the types of mixed feeling experience in scenarios such as bittersweetness ^26, 27^. Additionally, this previous study relied on univariate comparisons to make determinations about the uniqueness of mixed feeling states, which are likely to be less sensitive to differences in the spatiotemporal patterns.

In the present study, we had two main goals: 1. to determine if individualized feeling transitions during a dynamic, naturalistic experience can be predicted with fMRI and 2. to determine if various brain regions and networks differ in their representation of mixed feelings as a state with consistent and unique neural patterns. While in the scanner, the subjects watched the Academy Award nominated animated short *One Small Step* ^43^, a film that tells the story of a young girl who dreams of becoming an astronaut, and of her ever-supportive father. The film climaxes in a prototypical bittersweet moment when she achieves her goal yet reflects on the loss of her father.

After scanning, subjects rewatched the video and used buttons to label how they had felt when in the scanner. For our first goal, we used HMM as a data-driven approach to test if state transitions in a given neural region of interest would occur at similar time points to an individual’s emotion annotations. For the second, we used subjects’ individual emotion annotations as a ground truth to analyze whether mixed, positive, and negative states were neurally consistent and unique from each other.

## Results

Subjects reported strong variability in what they felt, when specific feelings occurred, and in the number of times their feelings changed (mean=12.5, standard deviation= 7.1, range=3-36).

### Matching individual’s feeling transitions to predicted neural state changes

In our first analysis, we tested whether HMM-predicted boundaries in a region of interest matched up with subjects’ reported feeling transitions at a rate that was significantly better than chance. For each subject, the model was fit so that k (the number of states for the data to be fit to) was equal to the number of feeling transitions reported by the subject plus 1, and a predicted boundary was considered a ‘match’ with subject report if within a 5-second window.

Significance was tested by permuting the timeseries and shuffling the boundary locations 10,000 times in each subject, for each region. Then, we aggregated each subjects’ distance from their null mean, and all the null distributions, to understand group-wide regional trends (Figure 2).

**Figure 1:**
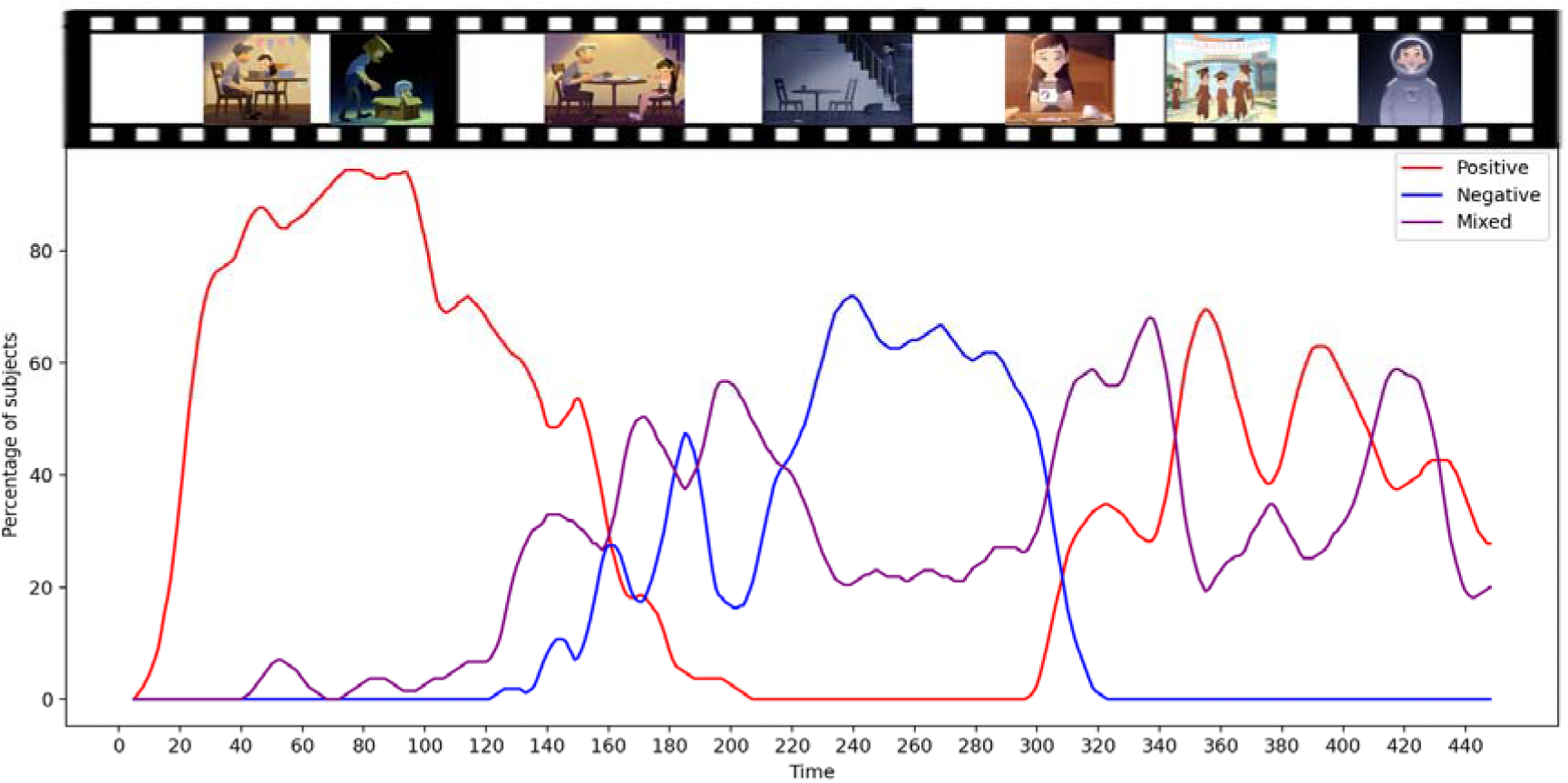
Percentage of subject’s feeling positive, negative, and mixed at each timepoint of the film. Percentage of subjects (N=27) reporting each state throughout the film.

**Figure 2:**
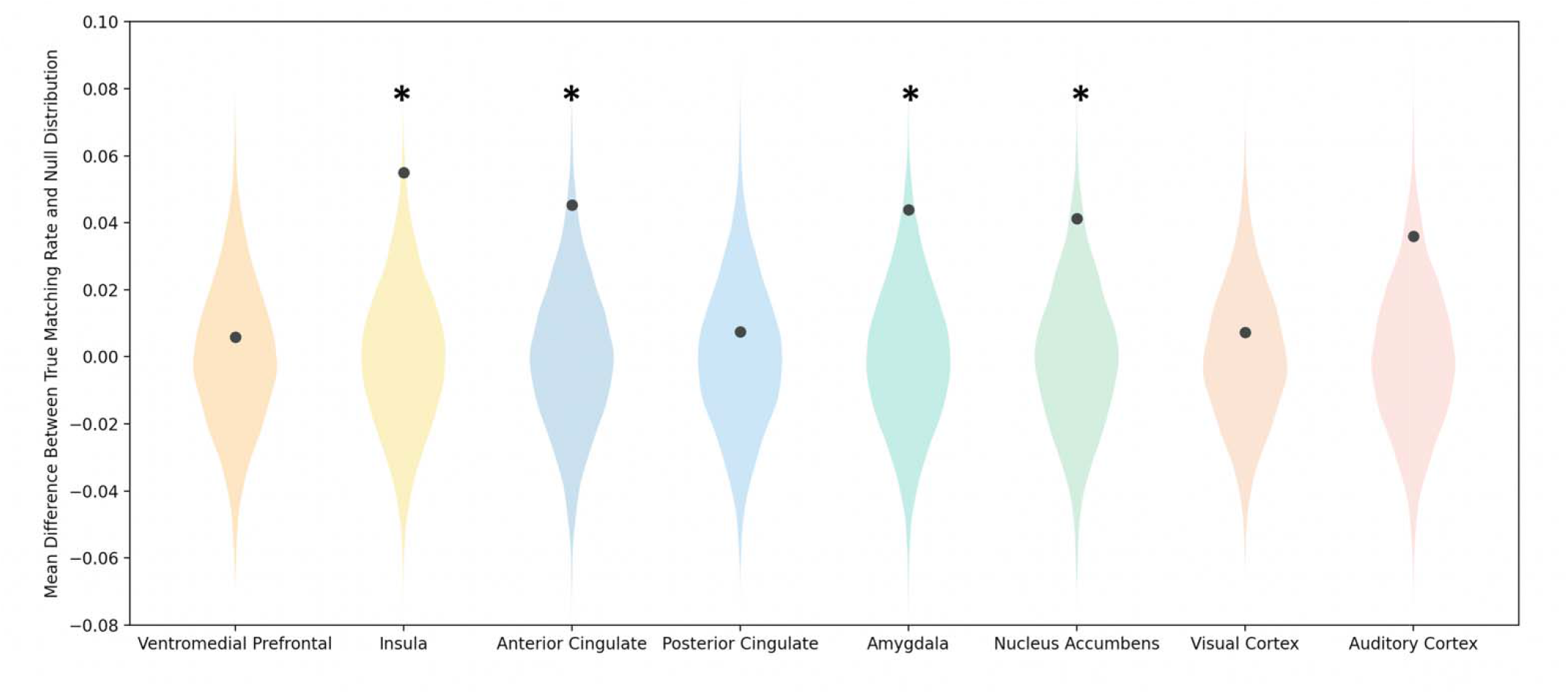
Significance of Regional Matching Rates Between Individual Self-Report and HMM Predicted Boundaries. Significance was determined by comparison with 10,000 permutations of shuffling TRs and boundaries. Each plot shows a distribution, across all subjects, of the average deviation from the mean of the null distribution for each individual subject. The dot indicates the true average difference between a subject’s matching rate and the mean of their null distribution. Asterisks indicates a matching rate significantly greater (p<0.05) than the null mean.

**Figure 3:**
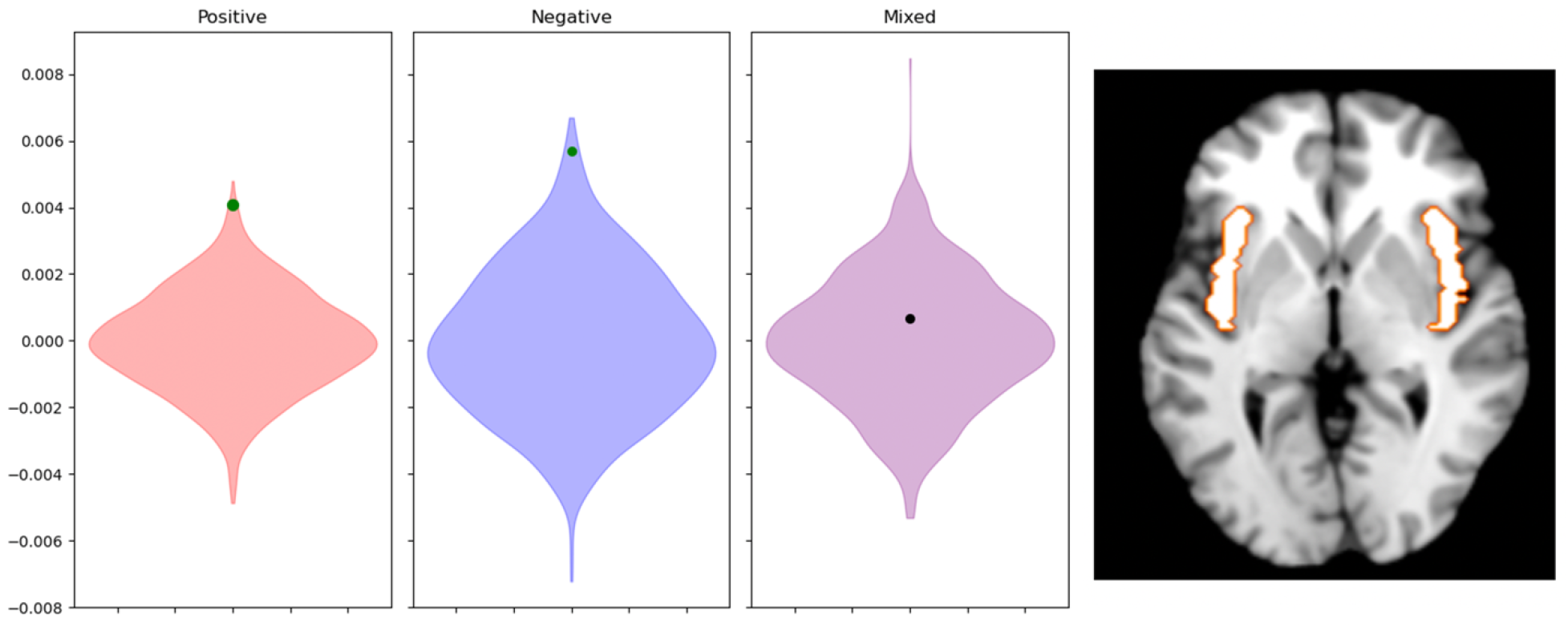
Average Difference in Neural Consistency from Null Distribution for Positive, Negative, and Mixed Valence Feelings in the Insula. Average difference between neural consistency of positive (red), negative (blue) and mixed (purple) timepoints across subjects, and the mean of individual subjects’ null distributions. Neural consistency is calculated as the average correlation between timepoints of the same feeling type subtracted by the average correlation of timepoints of that feeling type with other feeling types. Null distributions are made by 1000 permutations of shuffling TRs in each subject. Significance at p<0.05 is indicated by the true mean dot being green. In the top image, the dorsal anterior, ventral anterior, and posterior masks are show in orange, blue, and green respectively.

**Figure 4:**
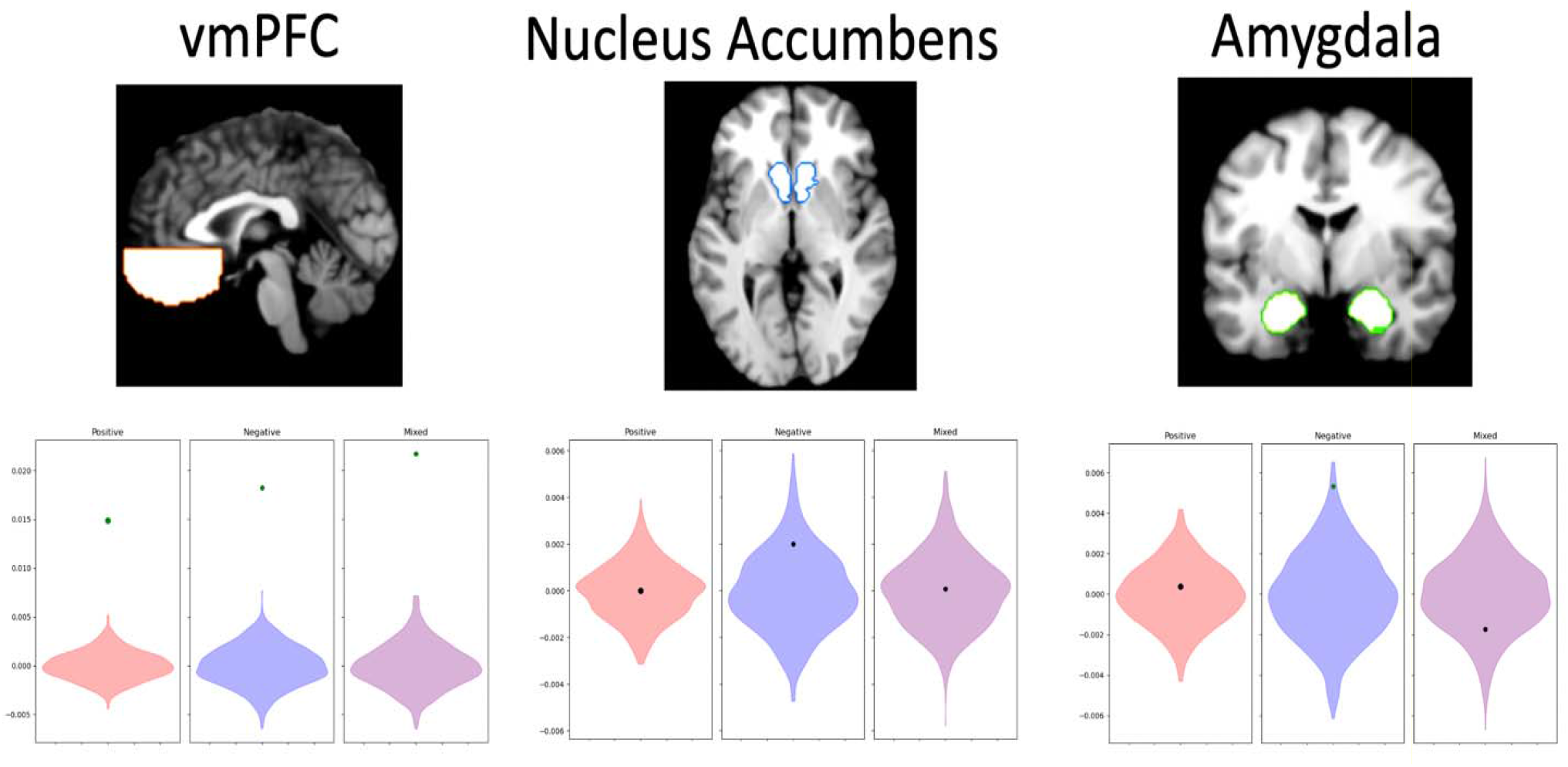
Average Difference in Neural Consistency from Null Distribution for Positive, Negative, and Mixed Valence Feelings in Ventromedial Prefrontal Cortex (vmPFC), Nucleus Accumbens, and Amygdala. Average difference between neural consistency of positive (red), negative (blue) and mixed (purple) timepoints across subjects, and the mean of individual subjects’ null distributions. Neural consistency is calculated as the average correlation between timepoints of the same feeling type subtracted by the average correlation of timepoints of that feeling type with other feeling types. Null distributions are made by 1000 permutations of shuffling TRs in each subject. Significance at p<0.05 is indicated by the true mean dot being green. Amygdala mask is shown in green and ventromedial prefrontal cortex mask is shown in orange.

**Figure 5:**
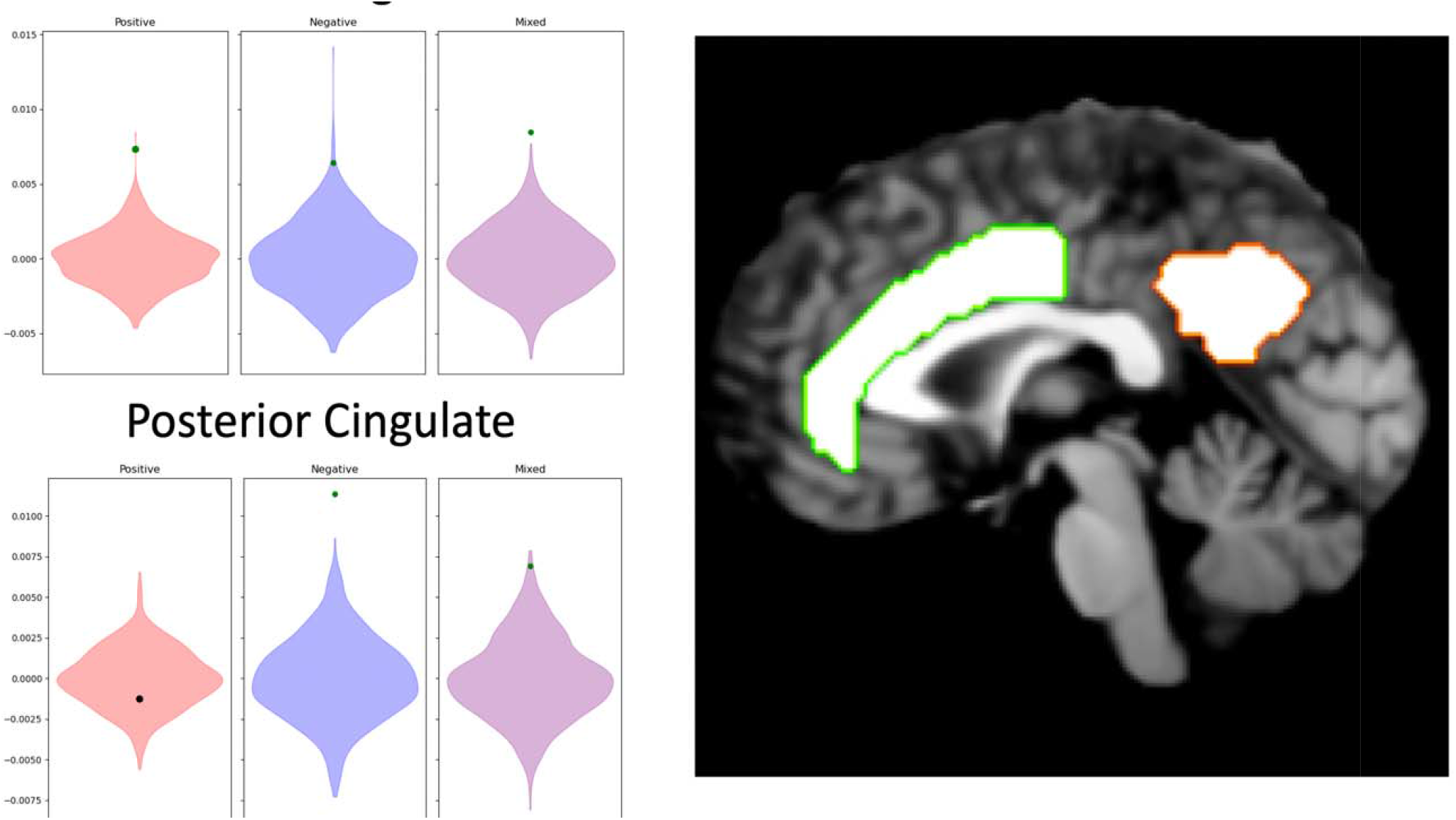
Average Difference in Neural Consistency from Null Distribution for Positive, Negative, and Mixed Valence Feelings in Cingulate Regions. Average difference between neural consistency of positive (red), negative (blue) and mixed (purple) timepoints across subjects, and the mean of individual subjects’ null distributions. Neural consistency is calculated as the average correlation between timepoints of the same feeling type subtracted by the average correlation of timepoints of that feeling type with other feeling types. Null distributions are made by 1000 permutations of shuffling TRs in each subject. Significance at p<0.05 is indicated by the true mean dot being green. Anterior cingulate mask is shown in green and the posterior cingulate mask is shown in orange.

**Figure 6:**
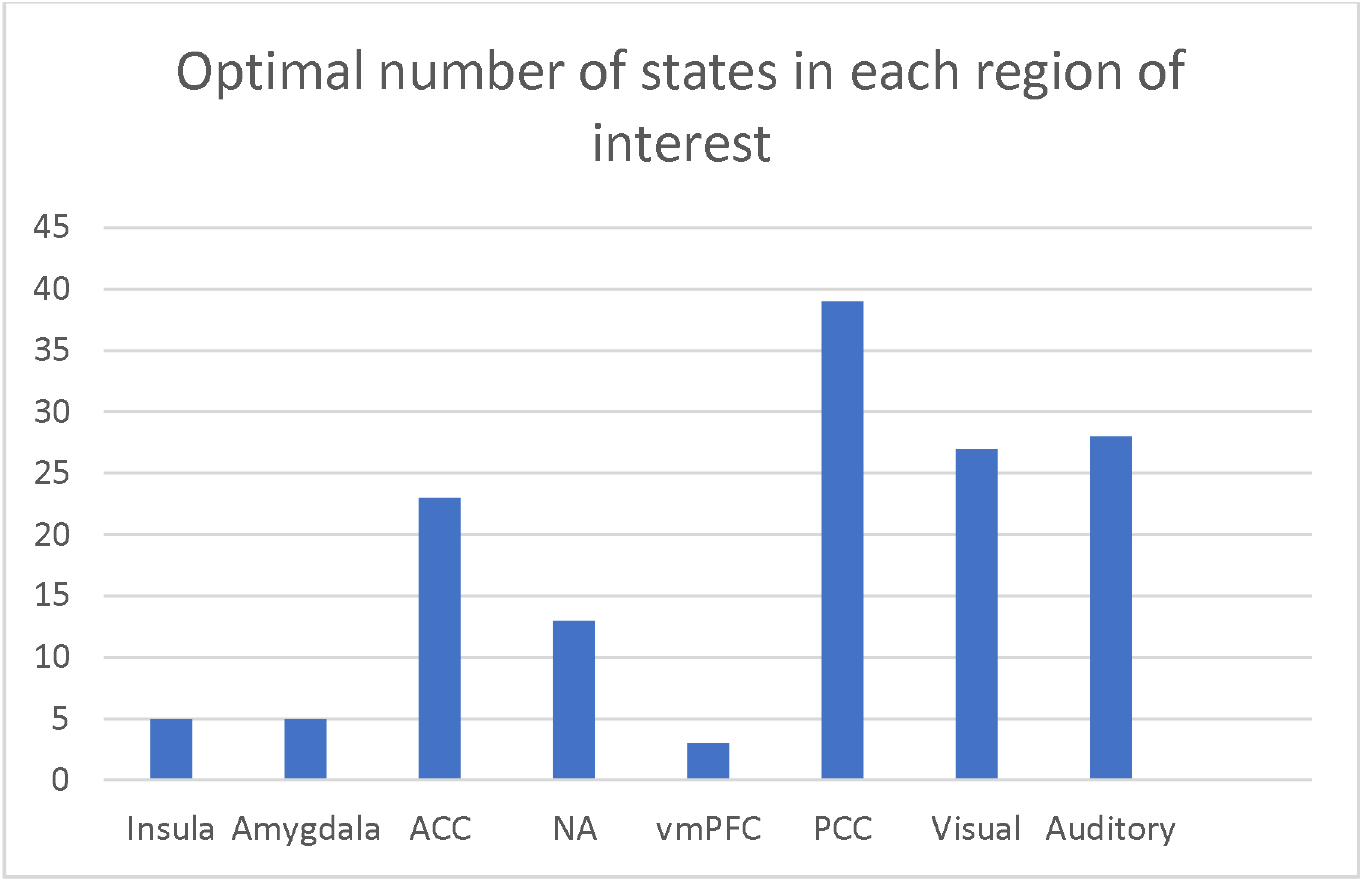
Optimal Number of States in Each Region Based on Within vs. Across Correlations in the Mean Data.

HMM-predicted boundaries matched subjects’ individual boundaries in the insula (33.2% match, p=0.007), amygdala (32.1% match, p=0.024), anterior cingulate (32.1% match, p=0.023), and nucleus accumbens (33.7% match, p=0.034) at rates significantly better than would be expected by chance. The ventromedial prefrontal cortex (28.3% match, p =0.383), posterior cingulate (27.0% match, p=0.357), visual cortex (25.2% match, p=0.354), and auditory cortex (31.2% match, p=0.056) did not significantly match with individuals’ self-report.

### Matching rates to individual vs consensus emotional report

We next tested the matching rate of each subject’s data with the average “consensus” emotion boundaries, an approach more similar to previous fMRI studies that employed HMMs ^11, 33, 44^. HMM-predicted boundaries did not significantly match consensus feeling transitions in any tested region.

Matching rates were significantly greater for individually reported feeling transitions vs. consensus feeling boundaries in all regions of interest: insula (match 33.2% vs 17.5%, p=0.0018), amygdala (32.1% vs. 13.8%, p=0.0017), anterior cingulate (32.1% vs. 14.8%, p=0.001), nucleus accumbens (33.7% vs. 17.5%, p=0.0023), posterior cingulate (27.0% vs 14.5%, p=0.005), ventromedial prefrontal cortex (28.3% vs. 15.3%, p=0.0032), auditory cortex (31.2% vs. 17.5%, p=0.0014), and visual cortex (25.2% vs. 13.8%, p=0.013).

### Matching rates to mean optimal regional boundaries

We additionally tested how boundaries derived from the mean neural data, across-subjects, and within each region, would fit with individual subjects’ data. This allowed us to explore how consistent predicted boundaries are in each of these regions when analysis decisions are agnostic to considerations of emotion. In sum: were the successful matching rates to individuals’ emotion report just reflective of regional consistencies in neural events overall? Based on the pattern of significant regions, this does not appear to be the case.

These mean-data derived boundaries significantly matched with boundaries predicted from individual subjects’ data in the insula (13.4% accuracy, p=0.046), ventral medial prefrontal cortex (42.1% match, p=0.01), anterior cingulate (40.7% match, p<0.0001), posterior cingulate (57.3% match, p<0.0001), visual cortex (74.4% match, p<0.0001), and auditory cortex (48.9% match, p<0.0001). Boundaries predicted from the mean of the group’s data notably did not match individually predicted boundaries in the amygdala (10.7% match, p=0.233) or nucleus accumbens (10.0% match, p=0.438).

### Neural consistency of positive, negative, and mixed valence feelings in different regions

In our final analysis, we aimed to test whether specific brain regions differed in the consistency of their neural states corresponding to positive, negative, and mixed feelings. We used each subjects’ feeling labels to determine periods of time where they reported feeling positively, negatively, and mixed. We then compared the difference in the average correlation of each timepoint of a feeling type with all other timepoints of that feeling type, with the average correlation of those timepoints correlated with all other timepoints that were of different feeling types (i.e. the correlation of all timepoints where a subject indicated mixed feelings with other timepoints where they reported mixed feelings, vs. the correlation of all mixed feeling timepoints with all positive and negative time points). In other words, we asked: at a timepoint labeled with a specific feeling type, is the pattern of voxels in a region more correlated with other timepoints that have the same feeling label than timepoints without that label? We will refer to this metric as *neural consistency*. Neural consistency for a specific state that is consistently above chance across subjects in a region demonstrates that 1) the spatial pattern of activity in that region is relatively consistent whenever it occurs and 2) that spatial pattern is significantly different from the other states it is being compared to. **A mixed state characterized by fluctuation, and/or similarity to the positive and negative states, would not pass this test.**

Analysis of the insular cortex found significant neural consistency for positive (p=0.002) and negative states (p=0.007), but not mixed (p=0.354). In the amygdala, only negative states (p=0.009) were significantly more neurally consistent than would be expected by chance (p=0.394 for positive; p=0.842 for mixed). The vmPFC was significant for all three feeling states (p<0.001 for all), as was the ACC (p=0.001 positive; p=0.007 negative; p<0.001 mixed). The PCC reached significance for negative (p<0.001) and mixed (p=0.004) but not for positive (p=0.767). In the auditory cortex all three reached significance (positive p<0.001, negative p=0.001, mixed p<0.001). In the visual cortex, all three reached significance (p<0.001). No feeling states were significantly consistent in the nucleus accumbens (p=0.524 positive, p=0.107 negative, p=0.472 mixed).

## Discussion

In this study, we used individualized feeling dynamics to understand the neural basis of mixed feelings. We focus on the relationship between individual affect and the individual’s data, rather than using consensus emotional features based on the stimuli. Overall, we found that brain state changes in the insular cortex, anterior cingulate, amygdala, and nucleus accumbens match significantly with feeling transitions that are defined by an individual’s self-report. All of these regions have been classically associated with interoception and salience processing, and represent crucial aspects of affective experience ^45, 46, 47, 48^. Furthermore, individually predicted boundaries matched significantly better with the subjects’ self-reported boundaries when compared to the emotion boundaries defined by consensus, which is the method use in most published analyses. While we often rely on assumed consistency of emotional features across subjects due to time constraints in collecting self-report data, and hoping to increase the power of the studies, defining stimuli in this manner likely shifts what our measures are studying further away from the neural correlates of conscious emotional experience and closer to other cognitive and stimuli-based features. An important advantage of naturalistic stimuli, when we have the corresponding self-report data available, is that we are encouraged to treat the “affect” in our study not just as a feature of the stimuli but as a feature that is also defined by the subject themselves ^12, 23^. It is possible that when we define emotion transitions based on consensus, the resulting boundaries correspond more closely to scene changes and narratively important moments in the film rather than to emotional idiosyncrasies between subjects. Therefore, these narrative understandings may be consistent even when affect is not.

Importantly, these findings cannot be explained by a general lack of consistency in any aspect of neurocognitive processing, or by the self-reported emotion transitions reflecting simply low-level stimulus features: our analysis of the optimal boundaries (derived from the mean data) revealed significant consistency in the timing of brain state changes across subjects, in many regions. These boundaries, however, 1) were not the same as the self-reported affect-related ones; 2) did not match at similar rates; and 3) significantly matched in many regions that did not significantly match with self-report, such as the ventromedial prefrontal cortex, visual cortex, and auditory cortex. Meanwhile the amygdala and nucleus accumbens, which significantly matched with subject self-report, did not have consistent optimal boundaries that fit consistently across all subjects.

Our analyses of the neural consistency of positive, negative, and mixed feelings fittingly had mixed results in terms of our initial hypotheses. We proposed that 1) since the insular cortex was involved in integrating our current affective moment it would have consistent neural states during mixed feelings as the key region involved in integrating the positive and negative inputs; 2) the subcortical structures, such as the amygdala and nucleus accumbens, that project to it would not have consistent neural states during mixed feelings; and 3) the anterior cingulate and prefrontal regions, to which the insular cortex projects, would fully flesh out the phenomenological experience of mixed feelings and thus have a neurally consistent state for it ^16^. The key deviation from our hypotheses was the finding that the insular cortex had consistent neural signatures for both positive and negative states, but not for mixed ones. The insular cortex has a role in integrating sources of affective information from multiple sources, and helps establish our subjective sense of the present moment ^38, 49, 50, 51^. We hypothesized that the temporal dynamics - on the scale of seconds in BOLD activity - would be consistent and unique in this region. Our reasoning for this hypothesis was that when integrating mixed valence, a unique pattern of activity compared to positive and negative valence would occur reflecting the integration process of the two types of valence information – while significant vacillation might occur across time on the neuronal level, we did not expect this pattern inconsistency to be reflected on the temporal scale of the BOLD signal might. It is instead possible that the neural fluctuations on the milliseconds scale are still reflected in a more in-flux BOLD response.

The anterior cingulate and ventromedial prefrontal cortex, the two regions we proposed as being crucial for elaborating on the experience of mixed feelings, had significant neural consistency during mixed feelings, as well as during positive and negative feelings. The juxtaposition between positive and negative feelings arising during the same situation often leads to a sense of conflict ^52, 53^. Activity in the anterior cingulate has long been linked to the experience of conflict and ambivalence, likely related to its proposed role in error monitoring ^54,55^. In the context of mixed feelings such as bittersweetness, the anterior cingulate may similarly allow the utilization of information about conflicting goals to be used in making complex decisions and self-regulation ^56, 57, 58^. The posterior cingulate also had a consistent neural signature for mixed feelings and negative feelings, however, it was not significantly neurally consistent for positive feelings, leaving room for the potential argument that the stability of the mixed feeling state is mostly driven by the component negativity. The posterior cingulate has been implicated in many studies of nostalgia as being important for the contextual and autobiographical processes involved in setting the stage for the bittersweetness of nostalgic remembrance ^58, 59^.

The ventromedial prefrontal cortex may play an important role in the representation of mixed feelings. Previous studies have found the orbitofrontal portion of the ventromedial prefrontal cortex to show unique activations in response to conflicting affective information ^60, 61,62^. More broadly, the ventromedial prefrontal cortex has been proposed to integrate affective information from the body with other sources of information, such as memories and conceptual knowledge ^39, 62, 63^. The integration of these various modalities may play a pivotal role in the generation of complex feelings. The one small-scale (n=10) fMRI study on reactions to bittersweet clips found a unique sub-region of higher activation in ventromedial prefrontal cortex when compared to positive and negative clips ^64^. The high distinctiveness of the three types of feelings in the ventromedial prefrontal cortex also aligns with the somatic marker hypothesis ^65^. Even if the various physiological patterns in insular cortex and subcortical regions are distinctively positive or negative, the vmPFC may facilitate the learning and representation of this fluctuating somatic pattern, leading to an emotional marker that is stable, yet representing mixed valence ^66, 67^.

Surprisingly, while all three feeling states were highly neurally consistent in the vmPFC, HMM methods could not predict the timings of emotion transitions at rates better than chance in this region. In the opposite vein, HMM significantly predicted emotion transitions in the nucleus accumbens despite the differently valenced states not being neurally consistent. It is possible that these findings hint at the temporal dynamics of these regions as well as the complexity of the information represented. Previous work has found that the rate of change in the vmPFC is relatively slow during naturalistic viewing, being on the order of 10s of seconds – but that these states themselves are quite stable^11, 34, 68^. The vmPFC’s slow and cumulative nature of information processing would mean that the changes between distinct states are more gradual, making it unlikely that boundaries could be predicted within a margin of error of 5 seconds as is done with our analyses. Meanwhile, subcortical regions such as the nucleus accumbens and amygdala, which are involved in quick saliency-related orienting, have dynamics characterized by briefer transitions in activity – new activity patterns may be more of ‘spikes’ than stable states^69^.

Sensory cortices also were also strongly neurally consistent for affective states, with the auditory cortex being significant for positive and mixed states, while the visual cortex was significant for positive, negative, and mixed states. Previous studies have found that the valence and emotion category of visuo-auditory stimuli can be predicted remarkably well from patterns of activity in lower-level sensory cortices ^70, 71, 72, 73^. Specific visual and auditory cues may have associations with specific types of affect. These types of features may additionally highlight a missing element of using media as ‘naturalistic’ stimuli. Recent reviews have pointed out that labeling media stimuli as ‘naturalistic’ may lead us to fail to consider the deliberate techniques used in media construction to induce specific experiences ^25, 74^. For sensory features qualities such as volume, brightness, etc, the inclusion of these features as regressors in neuroimaging analyses may help future studies understand how sensory features contribute to forming affective experience.

Beyond the sensory domain, these reviews also highlight that content analysis for why certain narrative and scene elements are designed the specific ways they are in pieces of media may help further parse out the specific cognitive processes involved in the induced affective experience. Future studies could investigate whether specific neural processes relate to the cognition involved in developing the narrative experience of the ‘bittersweet ending’ we used as a tool for inducing mixed feelings. Furthermore, it has been suggested that the poignancy and induction of mixed feelings in media is largely related to the ability to resonate with meaningful elements of the viewer’s life ^75^. The level of autobiographical relevance may modulate various neurocognitive processes, and could be investigated in future studies which integrate personal interviews.

## Conclusions

Our findings suggest that mixed feelings are associated with neurally consistent states in several cortical regions, indicating that they are not merely the result of a fluctuation in feeling or noise in the self-report. Furthermore, these results underscore the importance of considering individualized experiences when investigating the neural correlates of affect, because they may capture different and crucial elements of affective processing. Research on mixed feelings poses major difficulties, but the obstacles should not stand in the way of creating a comprehensive picture of the neurobiology of affect. The facts that mixed, positive, and negative feelings are unique and distinct from each other; and that they are consistent in anterior cingulate and ventromedial prefrontal cortex, suggest that their neurobiological features are **not** the mere consequence of switching back and forth between a positive and a negative state. The hypotheses related to whether the psychology of mixed feelings involves vacillation or a stable mixed state may never be fully falsifiable ^9^, but our data reveal a to a sufficiently stable neurobiological correlate for what had been defined via self-report. Our findings suggest that we should regard mixed valence as a valid and important concept.

## Methods

### Participants

28 right-handed, English-speaking subjects were recruited for the study. After one subject’s data was removed due to a scanning parameters error, the final sample included 27 right-handed, English-speaking subjects (mean age= 25.3, SD= 6.3; 15 female, 12 male) participated in the study. None had any history of neurological trauma. All provided informed consent as approved by the USC Institutional Review Board.

### Scanning parameters

All fMRI scanning was completed on a 3T Siemens Prisma System Scanner at the USC Dornsife Cognitive Neuroimaging Center using a 32-channel head coil. Anatomical images were acquired with a T1-weighted magnetization-prepared rapid gradient-echo (MPRAGE) sequence (repetition time [TR]/echo time [TE]=2300/2.26, voxel size 2 mm isotropic voxels, flip angle 9°). Functional images were acquired with a T2*-weighted gradient echo sequence (repetition time [TR]/echo time [TE]= 1000/35 ms, 41 transverse 3-mm slices, flip angle 52°, multiband factor=8). A T2-weighted volume was acquired for blind review by an independent neuroradiologist, in compliance with the scanning center’s policy and local IRB guidelines.

### Procedure

While in the scanner, subjects watched the Oscar-nominated animated short *One Small Step* (Chesworth & Pontillas, Taiko Studios, 2018). The film tells the story of a young girl who dreams of being an astronaut, and her shoe-making father who supports her dreams. The father encourages her dream as a child by making her astronaut boots, and when she gets older she begins to study astrophysics in college. She struggles in her courses and is initially rejected from the astronaut program. Throughout this struggle, every time she comes home her father is sitting at the kitchen table with food ready for her. One day, she returns home and he is not there- we then see her at his grave, crying. Later, when sorting through his belongings, she finds the old astronaut boot he had made for her. This rekindles her motivation; she begins to excel at school, and gets accepted to the astronaut program. We finally see her launch in a rocket ship to the moon, and when she takes the first step onto the surface, the scene pans to her as a child wearing the astronaut boots playing with her father on her bed.

After scanning, subjects re-watched the video and performed a feeling annotation task using a custom JavaScript application (https://github.com/jtkaplan/cinemotion) ). Subjects were instructed to reflect back on when they watched the video in the scanner, and to press buttons to indicate how they were feeling during that initial watching. Subjects were able to turn feeling labels on and off to indicate stretches of time where they felt “Positive”, “Negative”, and “Mixed”, and also had an additional button to indicate any period of time they had cried.

### fMRI preprocessing

Data were pre-processed using fMRIPrep ^76^. fMRIPrep implements brain extraction, slice-time correction, standard motion correction, spatial smoothing, and high-pass temporal filtering. Additionally ICA-AROMA was run for the removal of motion related components. For each preprocessed subject, we regressed out the effect of white matter, grey matter, and cerebrospinal fluid from the time course data. Finally, all subjects’ data were trimmed to be 454 TRs long, removing the opening 12 seconds of scanning and final 6 seconds. This helped align the neural data while accounting for both the initial 6 seconds delay before the video began, and 6 seconds of hemodynamic response function delay.

### Feeling labels

Each subject’s feeling labeling data was annotated to create individualized transition points for them. At each second of the clip, we determined which feeling state the subject was in based on the labels that were turned on. Positive and negative feelings were defined as moments where either label was on exclusively, while mixed feelings were defined as any moment the mixed feeling button was on, as subjects differed in whether they tended to use the mixed button as mutual exclusively to the other two, or tended to turn all three on together when experiencing mixed feelings. If no buttons were on, this was considered a neutral feeling period. The feeling labels were smoothed with a 5 second sliding window to account for mistaken button presses, small gaps of time between new labels, and delays between actual feeling onset and response time. Timepoints (to the nearest second) where feeling labels changed were then used as that subject’s specific transition points.

### Regions of interest (ROI)

In each of our analyses, we extracted voxel-wise timeseries from 8 regions of interest.

The insula ROI was obtained from merging the three functional connectivity based subdivisions in Deen et al, 2011 ^77^. The anterior cingulate, amygdala, and nucleus accumbens regions were obtained from the Harvard Cortical and Subcortical atlases. The posterior cingulate region was taken from Shirer et al., 2012. Finally, we used the planum temporale from the Harvard-Oxford cortical atlas as auditory cortex, Brodmann area 17 thresholded at 50% probability as early visual cortex.

### Matching individualized feeling boundaries to predicted neural state changes

All HMM models in this study were fitted using the BrainIak python package ^78^. The analysis fits an HMM model to the data by iteratively estimating the neural event signatures for a specific number of events (from here on referred to as k) and the temporal event structure, till the model converges on a high-likelihood solution ^33^.

For each subject, in each region of interest, we fit a separate HMM model. The K value for each subject was picked to be equal to the number of feeling transitions plus 1, so the model would fit the same number of state transitions as were reported subjectively while labeling the video.

We then compared the HMM-determined neural transition time points with that subject’s manually reported feeling transitions using a forward moving-window algorithm. The first neural boundary found within 5 TRs (TR=1 second) of the reported feeling boundary was considered a match. This resulted in an overall fractional match rate, representing the proportion of reported feeling boundaries that matched with neurally predicted boundaries for each region within each participant.

In testing for statistical significance, we aimed to account for the variability in the number of feelings subjects reported, and subjects’ respective neural differences. We first generated null distributions for every subject in every region by randomly shuffling the neural events, and randomly generating transition timepoints. We fit an HMM to these randomly shuffled data, and tested whether the randomly generated timepoints matched within 5 TRs of the HMM predicted ones, repeating this entire process 10,000 times. After the 10,000 permutations, we compared the subject’s true match rate to the mean of their null distribution to get a metric of how far their match rate was from chance. All these differences were then averaged across subjects to get one metric of the average difference between subjects’ match rates and their null match rates in a specific region. We repeated this process using all 10,000 permutations of each subject’s null distribution, resulting in an aggregated null distribution for the average difference metrics in each region. By comparing the actual average difference metrics to these aggregated null distributions, we were able to determine whether subjects’ neurally predicted boundaries were consistently matching with their reported boundaries across regions at a level that was significantly greater than chance.

### Matching consensus feeling boundaries to predicted neural state changes

We additionally tested the matching rate of each subject’s data with “consensus” emotion boundaries- an approach more similar to previous HMM fMRI studies ^11, 33, 44^. We defined consensus emotion boundaries as timepoints where 50% of subjects agreed on a new emotional state for at least 5 seconds. This led to a total of 7 boundaries across the length of the video. The HMM analysis, and null distributions, were then fit in the same manner as the analysis of individualized feeling boundaries. Using Wilcoxon signed-rank tests, we then compared the regional matching accuracy rates for individualized feelings vs. consensus boundaries.

### Consistent optimal boundaries in each region

We explored the optimal number of states in various regions, agnostic to any consideration of emotion. With the mean time-course data of all subjects, in each of our regions of interest, we ran HMM models with different k values ranging from 2-45. For each k-value (the number of states), we calculated the correlations of spatial signal patterns within predicted boundaries vs. across boundaries using the method originated in Baldassano, et al., 2017. Specifically, we calculated the correlation of every timepoint with the timepoint occurring 5 seconds later. The “within” correlation was determined as all of the timepoint correlations where both timepoints were within the boundaries of a predicted state. The “across” correlation was between timepoints outside a state and time points inside the state. If the period of time indicated by the state boundaries is a neurally consistent state in that region, timepoints within the state should be more spatially correlated with each other than with timepoints outside the state, despite being separated by the same temporal distance. Whichever k-value led to the greatest positive difference of the within correlation minus the across correlation was considered the optimal model for that region.

Using this optimal k value from the mean data, we then tested whether this same value, when applied to individual subjects’ data, would find similar boundary locations. In each region, for each subject, we used the best k value from the mean data in that region. We then compared the subject’s HMM determined transition points with the transition points from the mean data using a forward moving algorithm. The first neural boundary found within 5 TRs (TR=1 second) of the feeling boundary was considered a match. We calculated a matching accuracy rate for each subject in each region using this method.

To test for statistical significance, we first generated null distributions for every subject in every region by randomly shuffling the neural events, and randomly generating transition timepoints. We fit an HMM to these randomly shuffled data, and tested whether the randomly generated timepoints matched within 5 TRs of the HMM predicted ones, repeating this entire process 10,000 times. After the 10,000 permutations, we compared the subject’s true match rate to the mean of their null distribution to get a metric of how far their match rate was from chance. All these differences were then averaged across subjects to get one metric of the average difference between subjects’ match rates and their null match rates in a specific region. We repeated this process using all 10,000 permutations of each subject’s null distribution, resulting in an aggregated null distribution for the average difference metrics in each region. By comparing the actual average difference metrics to these aggregated null distributions, we were able to determine whether the differences between subjects’ neurally predicted boundaries were consistent across regions at a level that was significantly greater than chance.

The number of states that led to the highest within vs across correlation difference differed in each region. As expected, the optimal fit for sensory cortices involved a higher number of states than those for other regions, likely due to a greater number of changes in low-level sensory features in the video format. The k values displayed in Table 2 and Figure 2 were used in the next analysis.

**Table 2.**
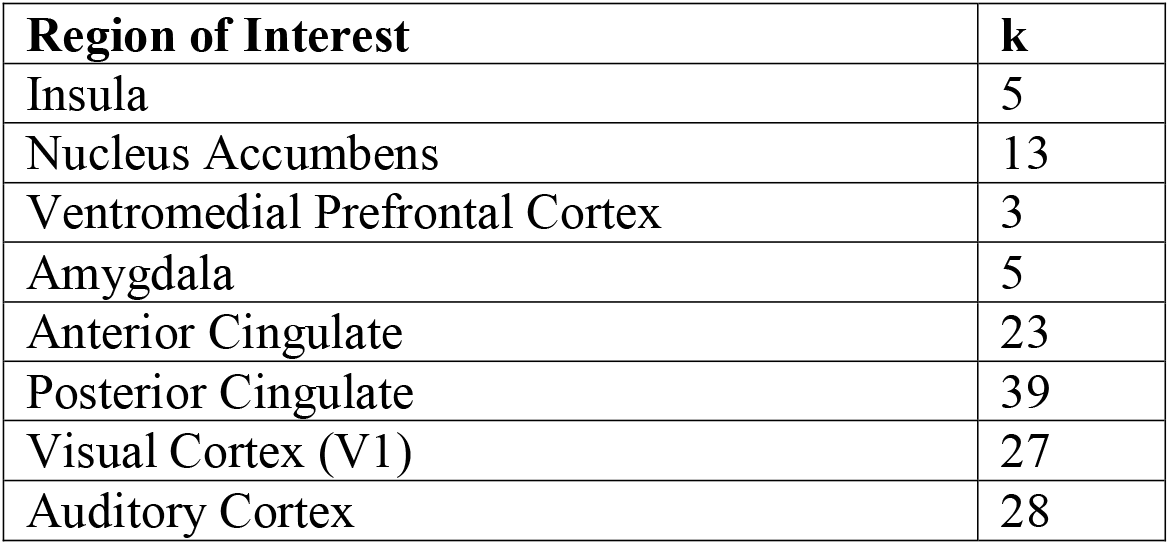

### Matching the regional optimal boundaries to individual subjects’ data

These mean-data derived boundaries significantly matched with boundaries predicted from individual subjects’ data in the insula (35% accuracy, p=0.046), ventral medial prefrontal cortex (42.1% match, p=0.01), anterior cingulate (10% match, p<0.0001), posterior cingulate (37.3% match, p<0.0001) visual cortex (33.4% match, p<0.0001), and auditory cortex (32.1% match, p<0.0001). Boundaries predicted from the mean of the group’s data notably did not match individually predicted boundaries in the amygdala (10% match, p=0.233) or nucleus accumbens (10% match, p=0.438).

### Neural consistency of feelings in different regions

In our final analysis, we aimed to test whether specific regions differed in the consistency of their neural states corresponding to positive, negative, and mixed feelings. To test this, we used each subjects’ feeling labels to determine periods of time where they reported feeling positively, negatively, and mixed. To quantify the consistency of neural states within subject-reported feeling state boundaries, we used a similar logic to the within vs. across correlation analysis for each feeling type. Specifically, we compared the difference in the average correlation of each timepoint of a feeling type with all other timepoints of that feeling type, with the average correlation of those timepoints correlated with all other timepoints that were of different feeling types (i.e. the correlation of all timepoints where a subject indicated mixed feelings with other timepoints where they reported mixed feelings, vs. the correlation of all mixed feeling timepoints with all positive and negative time points). We will refer to this metric as *neural consistency*. We specifically excluded timepoints where subjects did not report any feeling to avoid our results being driven largely by differences in how correlated affect timepoints are with neutral timepoints-our interest is in the distinguishability of different types of affective states. To create null distributions for each subject in each region, we shuffled the order of every TR 1000 times and calculated this correlation difference each time with the randomized TR’s. We then calculated a test-statistic by subtracting the subject’s true correlation difference with the mean of the null distribution. Finally, to create overall null distributions for each region, we used the same method as in the previous analyses: we calculated an average null test-statistic across all subjects for all 1000 of the permutations, and compared the true average test-statistic across the subjects to this distribution. Overall, this allowed us to test for an effect size of each feeling type being neurally more consistent, or less consistent, in each region of interest at a level that is higher than what would be expected by chance.

## References

1. Miyamoto Y, Uchida Y, Ellsworth PC. Culture and mixed emotions: co-occurrence of positive and negative emotions in Japan and the United States. Emotion 10, 404–415 (2010).

2. Lomas T. The value of ambivalent emotions: a cross-cultural lexical analysis. Qualitative Research in Psychology, 1–25 (2017).

3. Larsen JT, McGraw AP. The case for mixed emotions. Social and Personality Psychology Compass 8, 263–274 (2014).

4. Moeller J, Ivcevic Z, Brackett MA, White AE. Mixed Emotions: Network Analyses of Intra-Individual Co-Occurrences Within and Across Situations. Emotion, (2018).

5. Shirai M, Kimura T. Degree of Meaningfulness of an Event’s Ending Can Modulate Mixed Emotional Experiences Among Japanese Undergraduates. Perceptual and Motor Skills 129, 1137–1150 (2022).

6. Oliver MB, Woolley JK. Tragic and poignant entertainment: The gratifications of meaningfulness as emotional response. In: *The Routledge handbook of emotions and mass media*). Routledge (2010).

7. Abeyta AA, Routledge C. Nostalgia as a psychological resource for a meaningful life. The Happy Mind: Cognitive Contributions to Well-Being, 427–442 (2017).

8. Russell JA. Mixed emotions viewed from the psychological constructionist perspective. Emotion Review 9, 111–117 (2017).

9. Larsen JT. Holes in the Case for Mixed Emotions. Emotion Review 9, 118–123 (2017).

10. Barrett LF, Bliss-Moreau E. Affect as a Psychological Primitive. Adv Exp Soc Psychol 41, 167–218 (2009).

11. Chang LJ, et al. Endogenous variation in ventromedial prefrontal cortex state dynamics during naturalistic viewing reflects affective experience. Science Advances 7, eabf7129 (2021).

12. Adolphs R. How should neuroscience study emotions? by distinguishing emotion states, concepts, and experiences. Social Cognitive and Affective Neuroscience 12, 24–31 (2016).

13. Kragel PA, LaBar KS. Multivariate neural biomarkers of emotional states are categorically distinct. Soc Cogn Affect Neurosci 10, 1437–1448 (2015).

14. Nummenmaa L, Saarimaki H. Emotions as discrete patterns of systemic activity. Neurosci Lett, (2017).

15. Lench HC, Bench SW, Flores SA. Searching for evidence, not a war: Reply to Lindquist, Siegel, Quigley, and Barrett (2013).

16. Vaccaro AG, Kaplan JT, Damasio A. Bittersweet: The Neuroscience of Ambivalent Affect. Perspectives on Psychological Science 15, 1187–1199 (2020).

17. Berrios R, Totterdell P, Kellett S. Eliciting mixed emotions: a meta-analysis comparing models, types, and measures. Front Psychol 6, 428 (2015).

18. Kreibig SD, Gross JJ. Understanding Mixed Emotions: Paradigms and Measures. Curr Opin Behav Sci 15, 62–71 (2017).

19. Grossmann I, Ellsworth PC. What are mixed emotions and what conditions foster them? Life-span experiences, culture and social awareness. Current Opinion in Behavioral Sciences 15, 1–5 (2017).

20. Moore MM, Martin EA. Taking stock and moving forward: A personalized perspective on mixed emotions. Perspectives on Psychological Science 17, 1258–1275 (2022).

21. Rafaeli E, Rogers GM, Revelle W. Affective synchrony: individual differences in mixed emotions. Pers Soc Psychol Bull 33, 915–932 (2007).

22. Oh VY, Tong EM. Specificity in the Study of Mixed Emotions: A Theoretical Framework. Personality and Social Psychology Review 26, 283–314 (2022).

23. Saarimäki H. Naturalistic stimuli in affective neuroimaging: A review. Frontiers in human neuroscience 15, 675068 (2021).

24. Morgenroth E, Vilaclara L, Muszynski M, Gaviria J, Vuilleumier P, Van De Ville D. Probing Neurodynamics of Experienced Emotions-A Hitchhiker’s Guide to Film Fmri.

25. Grall C, Finn ES. Leveraging the power of media to drive cognition: A media-informed approach to naturalistic neuroscience. Social Cognitive and Affective Neuroscience 17, 598–608 (2022).

26. Janicke-Bowles SH, Bartsch A, Oliver MB, Raney AA. Traditional Conceptualizations of Eudaimonic Entertainment. The Oxford Handbook of Entertainment Theory, 363 (2021).

27. Ersner-Hershfield H, Mikels JA, Sullivan SJ, Carstensen LL. Poignancy: mixed emotional experience in the face of meaningful endings. Journal of personality and social psychology 94, 158 (2008).

28. Oliver MB. Tender affective states as predictors of entertainment preference. Journal of communication 58, 40–61 (2008).

29. Wulf T, Rieger D, Schmitt JB. Blissed by the past: Theorizing media-induced nostalgia as an audience response factor for entertainment and well-being. Poetics 69, 70–80 (2018).

30. Slater MD, Oliver MB, Appel M. Poignancy and mediated wisdom of experience: Narrative impacts on willingness to accept delayed rewards. Communication Research 46, 333–354 (2019).

31. Khoo GS. Contemplating tragedy raises gratifications and fosters self-acceptance. Human Communication Research 42, 269–291 (2016).

32. Greenwood D, Long CR. When movies matter: Emerging adults recall memorable movies. Journal of Adolescent Research 30, 625–650 (2015).

33. Baldassano C, Chen J, Zadbood A, Pillow JW, Hasson U, Norman KA. Discovering event structure in continuous narrative perception and memory. Neuron 95, 709–721. e705 (2017).

34. Lee CS, Aly M, Baldassano C. Anticipation of temporally structured events in the brain. Elife 10, e64972 (2021).

35. Yates TS, Skalaban LJ, Ellis CT, Bracher AJ, Baldassano C, Turk-Browne NB. Neural event segmentation of continuous experience in human infants. Proceedings of the National Academy of Sciences 119, e2200257119 (2022).

36. Tye KM. Neural Circuit Motifs in Valence Processing. Neuron 100, 436–452 (2018).

37. Berridge KC. Affective valence in the brain: modules or modes? Nature Reviews Neuroscience, (2019).

38. Craig AD. How do you feel--now? The anterior insula and human awareness. Nat Rev Neurosci 10, 59–70 (2009).

39. Barbas H. Connections underlying the synthesis of cognition, memory, and emotion in primate prefrontal cortices. Brain Res Bull 52, 319–330 (2000).

40. Wilson SJ, Creswell KG, Sayette MA, Fiez JA. Ambivalence about smoking and cue-elicited neural activity in quitting-motivated smokers faced with an opportunity to smoke. Addict Behav 38, 1541–1549 (2013).

41. Richard JM, Berridge KC. Nucleus accumbens dopamine/glutamate interaction switches modes to generate desire versus dread: D(1) alone for appetitive eating but D(1) and D(2) together for fear. J Neurosci 31, 12866–12879 (2011).

42. Murray RJ, Kreibig SD, Pehrs C, Vuilleumier P, Gross JJ, Samson AC. Mixed emotions to social situations: An fMRI investigation. NeuroImage 271, 119973 (2023).

43. Chesworth AP, B. One Small Step. Taiko Studios (2018).

44. Sachs ME, Ochsner K, Baldassano C. Brain state dynamics reflect emotion transitions induced by music. bioRxiv, (2023).

45. Damasio A, Carvalho GB. The nature of feelings: evolutionary and neurobiological origins. Nat Rev Neurosci 14, 143–152 (2013).

46. Seeley WW. The salience network: a neural system for perceiving and responding to homeostatic demands. Journal of Neuroscience 39, 9878–9882 (2019).

47. Critchley HD, Garfinkel SN. Interoception and emotion. Curr Opin Psychol 17, 7–14 (2017).

48. Saarimaki H, et al. Discrete Neural Signatures of Basic Emotions. Cereb Cortex 26, 2563–2573 (2016).

49. Picard F, Kurth F. Ictal alterations of consciousness during ecstatic seizures. Epilepsy Behav 30, 58–61 (2014).

50. Craig AD. Emotional moments across time: a possible neural basis for time perception in the anterior insula. Philosophical Transactions of the Royal Society B: Biological Sciences 364, 1933–1942 (2009).

51. Kent L, Wittmann M. Time consciousness: the missing link in theories of consciousness. *Neuroscience of Consciousness*, niab011 (2021).

52. Berrios R, Totterdell P, Kellett S. Investigating goal conflict as a source of mixed emotions. Cogn Emot 29, 755–763 (2015).

53. Mejia ST, Hooker K. Mixed Emotions Within the Context of Goal Pursuit. Curr Opin Behav Sci 15, 46–50 (2017).

54. Luttrell A, Stillman PE, Hasinski AE, Cunningham WA. Neural dissociations in attitude strength: Distinct regions of cingulate cortex track ambivalence and certainty. J Exp Psychol Gen 145, 419–433 (2016).

55. Cunningham WA, Raye CL, Johnson MK. Implicit and explicit evaluation: FMRI correlates of valence, emotional intensity, and control in the processing of attitudes. J Cogn Neurosci 16, 1717–1729 (2004).

56. Kruschwitz JD, et al. Anticipating the good and the bad: A study on the neural correlates of bivalent emotion anticipation and their malleability via attentional deployment. Neuroimage 183, 553–564 (2018).

57. Nohlen HU, van Harreveld F, Rotteveel M, Lelieveld GJ, Crone EA. Evaluating ambivalence: social-cognitive and affective brain regions associated with ambivalent decision-making. Soc Cogn Affect Neurosci 9, 924–931 (2014).

58. Yang Z, et al. Patterns of brain activity associated with nostalgia: a social-cognitive neuroscience perspective. Social Cognitive and Affective Neuroscience 17, 1131–1144 (2022).

59. Yang Z, Izuma K, Cai H. Nostalgia in the Brain. Current Opinion in Psychology, 101523 (2022).

60. Becker MP, et al. Altered emotional and BOLD responses to negative, positive and ambiguous performance feedback in OCD. Soc Cogn Affect Neurosci 9, 1127–1133 (2014).

61. Simmons A, Stein MB, Matthews SC, Feinstein JS, Paulus MP. Affective ambiguity for a group recruits ventromedial prefrontal cortex. Neuroimage 29, 655–661 (2006).

62. Rolls ET, Grabenhorst F. The orbitofrontal cortex and beyond: from affect to decision-making. Prog Neurobiol 86, 216–244 (2008).

63. Roy M, Shohamy D, Wager TD. Ventromedial prefrontal-subcortical systems and the generation of affective meaning. Trends in cognitive sciences 16, 147–156 (2012).

64. Schulte FP, Maderwald S, Kraemer NC, Brand M. Bittersweet-neither happy nor sad. An experimental comparison of the neural effects of bittersweet, negative, and positive film clips using 7T fMRI. Proc Intl Soc Mag Reson Med 20, 2136 (2012).

65. Damasio AR. The somatic marker hypothesis and the possible functions of the prefrontal cortex. Philos Trans R Soc Lond B Biol Sci 351, 1413–1420 (1996).

66. Panksepp J. On the embodied neural nature of core emotional affects. Journal of consciousness studies 12, 158–184 (2005).

67. Dunning D, Fetchenhauer D, Schlösser T. The varying roles played by emotion in economic decision making. Current opinion in behavioral sciences 15, 33–38 (2017).

68. Honey CJ, et al. Slow cortical dynamics and the accumulation of information over long timescales. Neuron 76, 423–434 (2012).

69. Puccetti NA, Villano WJ, Fadok JP, Heller AS. Temporal dynamics of affect in the brain: Evidence from human imaging and animal models. Neuroscience & Biobehavioral Reviews 133, 104491 (2022).

70. Kragel PA, Reddan MC, LaBar KS, Wager TD. Emotion schemas are embedded in the human visual system. Science advances 5, eaaw4358 (2019).

71. Sachs ME, Habibi A, Damasio A, Kaplan J. Decoding the neural signatures of emotions expressed through sound. NeuroImage 174, 1–10 (2018).

72. Bo K, et al. Decoding neural representations of affective scenes in retinotopic visual cortex. Cerebral Cortex 31, 3047–3063 (2021).

73. Ethofer T, Van De Ville D, Scherer K, Vuilleumier P. Decoding of emotional information in voice-sensitive cortices. Current biology 19, 1028–1033 (2009).

74. Schmaelzle R, Huskey R. Integrating Media Content Analysis, Reception Analysis, and Media Effects Studies. Frontiers in Neuroscience 17, 612 (2023).

75. Klimmt C, Rieger D. Biographic resonance theory of eudaimonic media entertainment. The Oxford handbook of entertainment theory, 383–402 (2021).

76. Esteban O, et al. fMRIPrep: a robust preprocessing pipeline for functional MRI. Nature methods 16, 111–116 (2019).

77. Deen B, Pitskel NB, Pelphrey KA. Three systems of insular functional connectivity identified with cluster analysis. Cerebral cortex 21, 1498–1506 (2011).

78. Kumar M, et al. BrainIAK: The brain imaging analysis kit. Aperture neuro 1, (2021).

